# Integrated multi-omic analysis reveals novel subtype-specific regulatory interactions in pediatric B-cell acute lymphoblastic leukemia

**DOI:** 10.1101/2025.08.13.670107

**Authors:** Irina Pushel, Zachary S Clark, Lisa A Lansdon, Byunggil Yoo, Michaella J Rekowski, Nicole M Wood, Michael P Washburn, Midhat S Farooqi

## Abstract

Molecular subtyping of pediatric B-cell acute lymphoblastic leukemia (B-ALL) has improved patient outcomes through stratification and selection of targeted therapies. Despite extensive genomic and transcriptomic profiling of this cancer, few studies to date have characterized the proteomic landscape, although proteins are the direct targets of many therapeutic agents. In this study, we demonstrate the utility of multi-omic integration of global transcriptomic, proteomic, and phosphoproteomic profiles of samples from patients diagnosed with either of two B-ALL subtypes – Ph- like (*BCR::ABL1*-like) and *ETV6::RUNX1*. Through individual and multi-omic analysis, we recapitulate known transcriptomic findings and identify novel subtype-specific proteomic and phosphoproteomic biomarkers. Our findings suggest a previously undescribed role for calcium-dependent signaling processes in Ph-like B-ALL, which has the potential to serve as a novel avenue for targeted treatments. By integrating multiple ‘omics modalities, we identify not only features of interest but also begin to unravel the regulatory interactions driving subtype-specific mechanisms of leukemogenesis. This integrated analytic approach paves the way for enhanced precision medicine for precise subtyping and treatment selection for pediatric leukemia patients. Mass spectrometry data generated in this study have been deposited in MassIVE under accession MSV000097955.

## Introduction

B-cell acute lymphoblastic leukemia (B-ALL) is one of the most common childhood cancers, accounting for nearly a quarter of pediatric cancer diagnoses. Genetic subtyping of B-ALL has improved patient stratification and the selection of targeted therapies, thereby improving outcomes (J. Li et al., 2021; Tasian et al., 2015). Although subtyping has largely been focused on DNA lesions known to be cancer drivers, additional data, including gene expression profiling, is beginning to be integrated into clinical diagnostic practice (Hu et al., 2023; Vaske et al., 2019).

While genomic and transcriptional profiling have provided valuable insights into the molecular processes driving individual leukemic subtypes, these approaches are limited in that they do not consider cell state regulation beyond transcription. In recent years, proteomic and phosphoproteomic approaches have shed light on signaling pathways driving cancers and have facilitated the identification of targeted treatment options (Casado et al., 2018, 2023; Nguyen et al., 2021; Veltri et al., 2022). However, to date, efforts to integrate proteomic and phosphoproteomic characterization to leukemic subtyping have been limited (Jayavelu et al., 2022; Kourti et al., 2023). To extend these approaches to pediatric leukemias, we profile samples from patients diagnosed with one of two well-defined subtypes: *BCR::ABL1-*like and *ETV6::RUNX1* B-ALL, which have distinct prognoses, known signaling pathways, and treatment approaches.

*BCR::ABL1*-like (also known as Philadelphia-like, or Ph-like) B-ALL is an aggressive leukemic subtype and is diagnosed in approximately 10-20% of pediatric B-ALL patients (Bernt & Hunger, 2014). As the name suggests, this subtype is characterized by a transcriptional profile similar to the equally aggressive *BCR::ABL1* (Ph+) B-ALL, but lacks the signature gene fusion. The transcriptional similarity between these subtypes underlies the activation of kinase signaling, driving the leukemic phenotype (Harvey et al., 2010; Mullighan et al., 2009; Roberts et al., 2012; Schmäh et al., 2017). Recent studies have shown improved outcomes for pediatric Ph-like B-ALL patients treated with tyrosine kinase inhibitors (TKIs), leveraging these mechanistic findings (Kaczmarska et al., 2021; Reshmi et al., 2017; Roberts et al., 2014a).

In contrast, *ETV6::RUNX1* B-ALL is the most common subtype of B-ALL, characterized by the fusion of *ETV6* and *RUNX1* genes. It has a favorable prognosis, and de-escalation of chemotherapy intensity in patients with this rearrangement is currently being investigated (Østergaard et al., 2024). Prior work suggests that the *ETV6::RUNX1* fusion is the first hit of a “two-hit” leukemia, with a second lesion (such as loss of *ETV6* or gain of *RUNX1*, among others) acting as the driver of the leukemia (Kaczmarska et al., 2023; Rodríguez-Hernández et al., 2021). Considering the differences in prognosis, treatment response, and known activation of kinase signaling in Ph-like B-ALL versus *ETV6*::*RUNX1* B- ALL, we felt that a comparison of these subtypes would serve as a strong proof of concept to explore the value of a multi-omic approach to characterizing subtype-specific mechanisms.

In this study, we performed proteomic and phosphoproteomic profiling of samples collected at diagnosis and remission from pediatric leukemia patients with one of two subtypes of B-ALL: *BCR::ABL1-*like (Ph-like) and *ETV6::RUNX1*. We analyzed each dataset, as well as existing RNAseq data at diagnosis, to identify subtype-specific features, which revealed increased calcium-dependent signaling in Ph-like samples. We then performed an integrated analysis of all three datasets, which enabled us to identify multiple layers of regulation, including subtype-specific phosphorylation of known cancer-associated proteins such as IGF2BP1, MS4A1, and BCLAF1. Taken together, these data add to our understanding of the molecular profile of Ph-like and *ETV6::RUNX1* B-ALL, demonstrating the utility of multi-omic comparison in pediatric leukemia subtype characterization.

## Materials and Methods

### Patient samples

For each subtype, 5 patients were chosen with frozen blood and/or bone marrow aspirate (at least 2×10^6^ cells) collected at diagnosis and remission within the Children’s Mercy Research Institute Biorepository (CRIB). This research was conducted in accordance with the Declaration of Helsinki and approved by the Children’s Mercy Institutional Review Board. All patient samples were collected with written informed consent of the parents/guardians and assent of patients.

### Proteomics and phosphoproteomics

Cells were lysed by resuspending in 100 µl of RIPA buffer with protease and phosphatase inhibitors and nuclease per 1×10^6^ cells and incubating on ice for 30 minutes followed by sonicating in a water bath for 15 minutes. Samples were centrifuged at 14000 *x* g for 10 minutes at 4 °C and lysates were transferred to new tubes. Samples were reduced with the addition of 0.5 M TCEP to a final concentration of 5 mM followed by incubation at 37°C for 30 minutes. Reduced samples were alkylated with the addition of 375 mM iodoacetamide to a final concentration of 10 mM followed by incubation in the dark at room temperature for 30 minutes. Ice cold acetone was added to each sample to a volume ratio of 5:1. Samples were vortexed and stored at -20°C overnight. After precipitation, samples were centrifuged at 14,000 x *g* at 4°C for 30 minutes to pellet the proteins. The supernatant was removed, and the pellet was air dried on benchtop for 10 minutes. The proteins were resuspended in 50 mM TEAB pH 8, 2 mM CaCl2. Trypsin was added (500 ng) and the proteins were allowed to digest overnight at 37°C with shaking at 500 RPM (Thermomixer, Eppendorf). The digestion was quenched with the addition of 10% formic acid to a final concentration of 1%. The peptides were enriched for phosphorylation by the SMOAC method (Choi et al., 2017). The peptides that did not bind the resin were analyzed as the global sample and the enriched samples were pooled and run as the phosphopeptide enriched fraction. Peptide concentration was measured using a Nanodrop spectrophotometer (Thermo Scientific) at 205 nm prior to LC-MS/MS analysis.

Samples were injected using the Vanquish Neo (Thermo) nano-UPLC onto a C18 trap column (PepMap™ Neo Trap, 0.3 mm x 5 mm, 5 µm particle size) using pressure loading. Peptides were eluted onto the separation column (PepMap™ Neo, 75 µm x 150 mm, 2 µm C18 particle size, Thermo) prior to elution directly to the mass spectrometer. Briefly, peptides were loaded and washed for 5 minutes at a flow rate of 0.350 µL/min at 2% B (mobile phase A: 0.1% formic acid in water, mobile phase B: 80% ACN, 0.1% formic acid in water). Peptides were eluted over 100 minutes from 2-25% mobile phase B before ramping to 40% B in 20 min. The column was washed for 15 min at 100% B before re-equilibrating at 2% B for the next injection. The nano-LC was directly interfaced with the Orbitrap Ascend Tribrid mass spectrometer (Thermo) using a silica emitter (20 µm i.d., 10 cm, CoAnn) equipped with a high field asymmetric ion mobility spectrometry (FAIMS) source. The data were collected by data dependent acquisition with the intact peptide detected in the Orbitrap at 120,000 resolving power from 375-1500 *m/z*. Peptides with charge +2-7 were selected for fragmentation by higher energy collision dissociation (HCD) at 28% NCE and were detected in the ion trap at rapid scan rate (global) or in the Orbitrap at 30,000 resolving power (enriched). Dynamic exclusion was set to 60s after one instance. The mass list was shared between the FAIMS compensation voltages. FAIMS voltages were set at -45 (1.4 s), -60 (1 s), -75 (0.6 s) CV for a total duty cycle time of 3s. Source ionization was set at +1700 V with the ion transfer tube temperate set at 305 °C. Raw files were searched against the human protein database downloaded from Uniprot on 05-05-2023 and a common contaminants database with variable phosphorylation allowed on S, T, and Y residues (enriched) using SEQUEST in Proteome Discoverer 3.0 (Orsburn, 2021). Abundances, abundance ratios, and p-values were exported to Microsoft Excel for further analysis with packages available in R.

### Downstream data analysis and differential expression

Downstream analysis and visualization were performed in R 4.3.3. Protein abundance normalization was performed using proDA 1.16.0 (Ahlmann-Eltze & Anders, 2020). Pathway enrichment was performed using gProfiler2 0.2.3 (Kolberg et al., 2023). UpSet plots generated using UpSetR 1.4.0 (Conway et al., 2017). Kinase activity inferred using KSEA App 1.0 (Wiredja et al., 2017).

### RNA sequencing and analysis

Patient blood and/or bone marrow samples from the CRIB had RNA isolated and libraries prepared with the Illumina Stranded Total RNA with RiboZero kit. Paired end sequencing was performed on an Illumina NovaSeq 6000. Reads were aligned to GRCh38 using STAR 2.6.0 (Dobin et al., 2012) and read counts estimated using kallisto 0.46.2 (Bray et al., 2016). Raw and processed data are available through the Childhood Cancer Data Initiative at https://datacatalog.ccdi.cancer.gov/dataset/CCDI-phs002529. Differential expression was performed using DESeq2 1.42.1 (Love et al., 2014) in R 4.3.3. Visualization and pathway enrichment performed as described above.

### Integrative multi-omic analysis

omicade4 1.42.0 (Meng et al., 2014) was used for hierarchical clustering of individual assays and integrated analysis across transcriptomic, proteomic, and phosphoproteomic datasets. Multiple co-inertia analysis (MCIA) was performed on nine samples (all (phospho)proteomics samples excluding Ph4) with 8 axes kept in the analysis (cia.nf). Pseudo-eigenvalues, representing contributions of each assay to dimensions 1-2, were extracted from the mcoin$mcoa$lambda object. Subtype-specific drivers from all assays were selected using the selectVar function on dimension 2, with bounds of (1, Inf) and (- Inf, -1). Network visualization was performed using Cytoscape 3.10.3 (Shannon et al., 2003) – log2(fold change) of Ph-like vs. *ETV6::RUNX1* samples at diagnosis, as well as predicted kinase enrichment (activity) from KSEA App were used as inputs. Phosphosite targets from KSEA App output were used to establish the network, with each phosphosite (phosphorylation event) treated as an individual arrow.

### Proteomic and phosphoproteomic data availability

The mass spectrometry data have been deposited to MassIVE. The accession number for the data reported in this paper is MassIVE MSV000097955. It has also been submitted to ProteomeXchange (PXD064162).

### Code availability

Code for the analysis performed in this paper is available at https://github.com/ChildrensMercyResearchInstitute/b-all_proteomics2025

## Results

### Establishing a framework for proteomic characterization of pediatric B-ALL

To understand the molecular drivers of distinct subtypes of pediatric B-ALL, we incorporated existing genomic and transcriptomic data with newly generated global proteomic and phosphoproteomic profiles for a cohort of patients with either *ETV6::RUNX1* B-ALL or Ph-like B-ALL. We identified five patients for each subtype with samples collected at both diagnosis and remission available in the CRIB for proteomic and phosphoproteomic data generation (Table 1). When possible, we selected patients with existing genomic and/or transcriptomic data.

**Table 1.** Patient characteristics. Genomic findings and corresponding assays listed for each patient.

| Patient ID | Patient ID2 | Subtype | Genomic findings | Assay(s) |
| --- | --- | --- | --- | --- |
| TB-000018 | ER1 | *ETV6::RUNX1* | *ETV6::RUNX1*, loss of *ETV6* (clonal) | Chromosomes, FISH, Microarray |
| TB-000023 | ER2 | *ETV6::RUNX1* | *ETV6::RUNX1*, gain of *ETV6::RUNX1* (subclonal), loss of *ETV6* (subclonal) | Chromosomes, FISH, Microarray |
| TB-000025 | ER3 | *ETV6::RUNX1* | *ETV6::RUNX1*, loss of *ETV6* (subclonal) | Chromosomes, FISH, Microarray |
| TB-000042 | ER4 | *ETV6::RUNX1* | *ETV6::RUNX1*, loss of *PAX5* exons 2-5 | Chromosomes, FISH, Microarray |
| TB-000203 | ER5 | *ETV6::RUNX1* | *ETV6::RUNX1*, loss of last 3 exons of *ETV6* | Chromosomes, FISH, Microarray |
| TB-000005 | Ph1 | Ph-like | *P2RY8::CRLF2* | Chromosomes, FISH, Microarray |
| TB-000071 | Ph2 | Ph-like | *IGH::CRLF2* | Chromosomes, FISH, UCSF500 (NGS) |
| TB-000115 | Ph3 | Ph-like | unknown | No diagnostic drivers found via Chromosomes, FISH, Microarray, UCSF500 (NGS). TriCore LDA positive for Ph-like expression, high CRLF2 expression, negative for fusions |
| TB-000143 | Ph4 | Ph-like | *P2RY8::CRLF2* | Chromosomes, FISH, Microarray |
| TB-000174 | Ph5 | Ph-like | *RCSD1::ABL2* | Chromosomes, FISH, Nationwide Archer FusionPlex |

We analyzed individual ‘omics data separately using current standard approaches, then performed an integrated analysis to identify subtype-specific regulatory mechanisms (Figure 1). Importantly, even patients with the same B-ALL subtype showed variability in genomic drivers, reflecting the highly individualized nature of cancer and underscoring the value of personalized ‘omics analysis in pediatric leukemia diagnosis and treatment selection.

**Figure 1.**
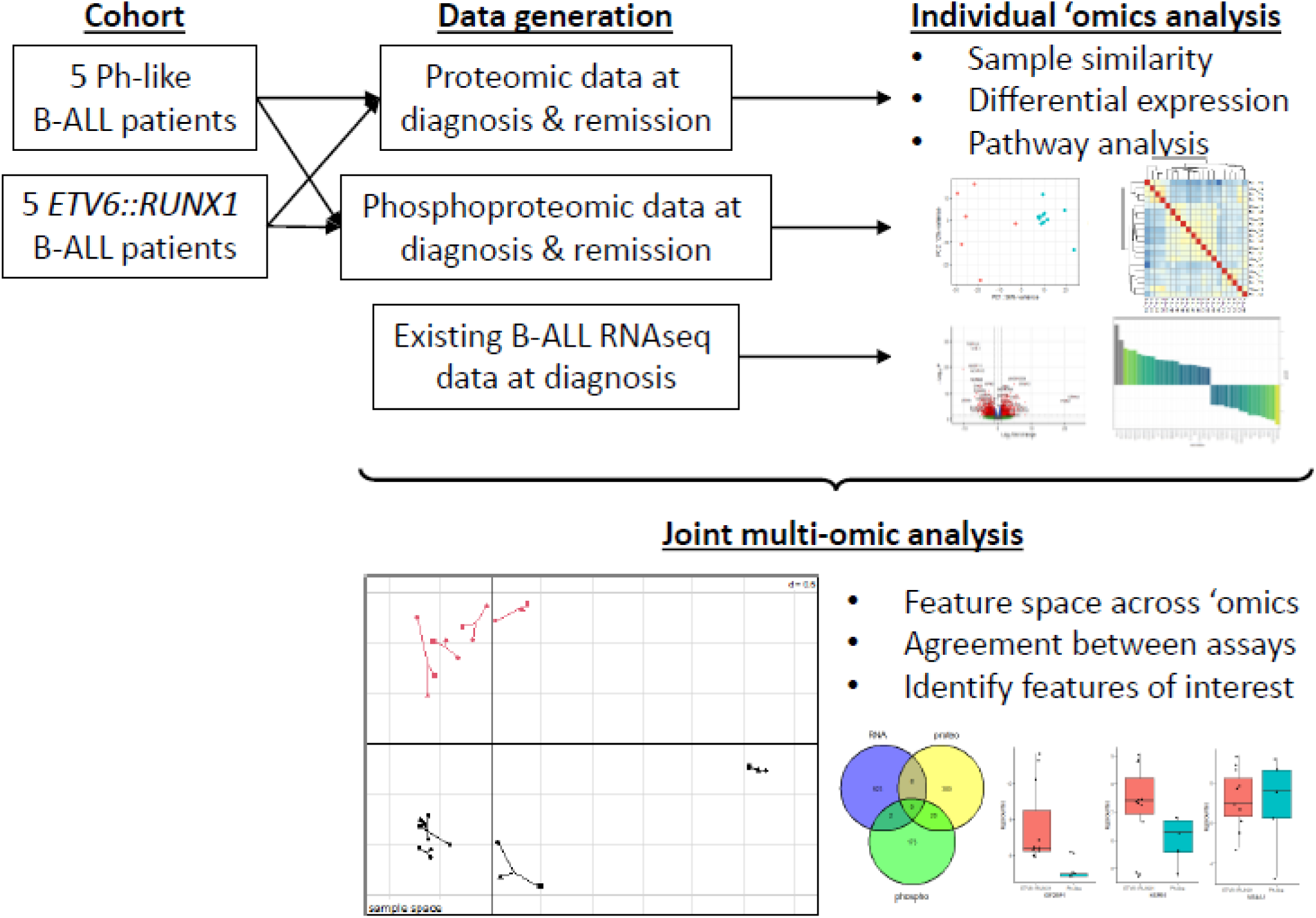
Schematic of study. Five patients each with Ph-like and *ETV6::RUNX1* B-ALL were selected from the Children’s Mercy Research Institute Biorepository for proteomic and phosphoproteomic analysis. Together with existing RNAseq data, individual ‘omics analysis was performed to identify subtype-specific insights, followed by integrative multi-omic analysis to extract commonalities and regulatory relationships between features.

For each sample, we generated global proteomic and phosphoproteomic profiles using Data Dependent Acquisition (DDA). In proteomic data, samples collected at remission tended to show higher similarity to each other regardless of subtype, while samples collected at diagnosis showed subtype-specific similarity (Figure 2A). Following principal component analysis (PCA), we saw that individual patients showed consistent separation of diagnosis (right) and remission (left) samples across dimension 1 (Figure 2B). Interestingly, while dimension 2 did not show separation based on any known factors of interest, we did see separation based on subtype across dimension 3, which accounts for nearly as much of the variance in the dataset as dimension 2 (Supplementary Figure 1A). Based on this observation, we extracted the top five proteins driving separation across dimensions 1 and 3, which included NDUFA6, SNRPD1, ATP5MG, ETFA, and HINT2 driving diagnosis/remission differences and TM9SF3, JCHAIN, NEIL1, ORM1, and SPAG17 driving subtype-specific differences (Supplementary Figure 1B).

**Figure 2.**
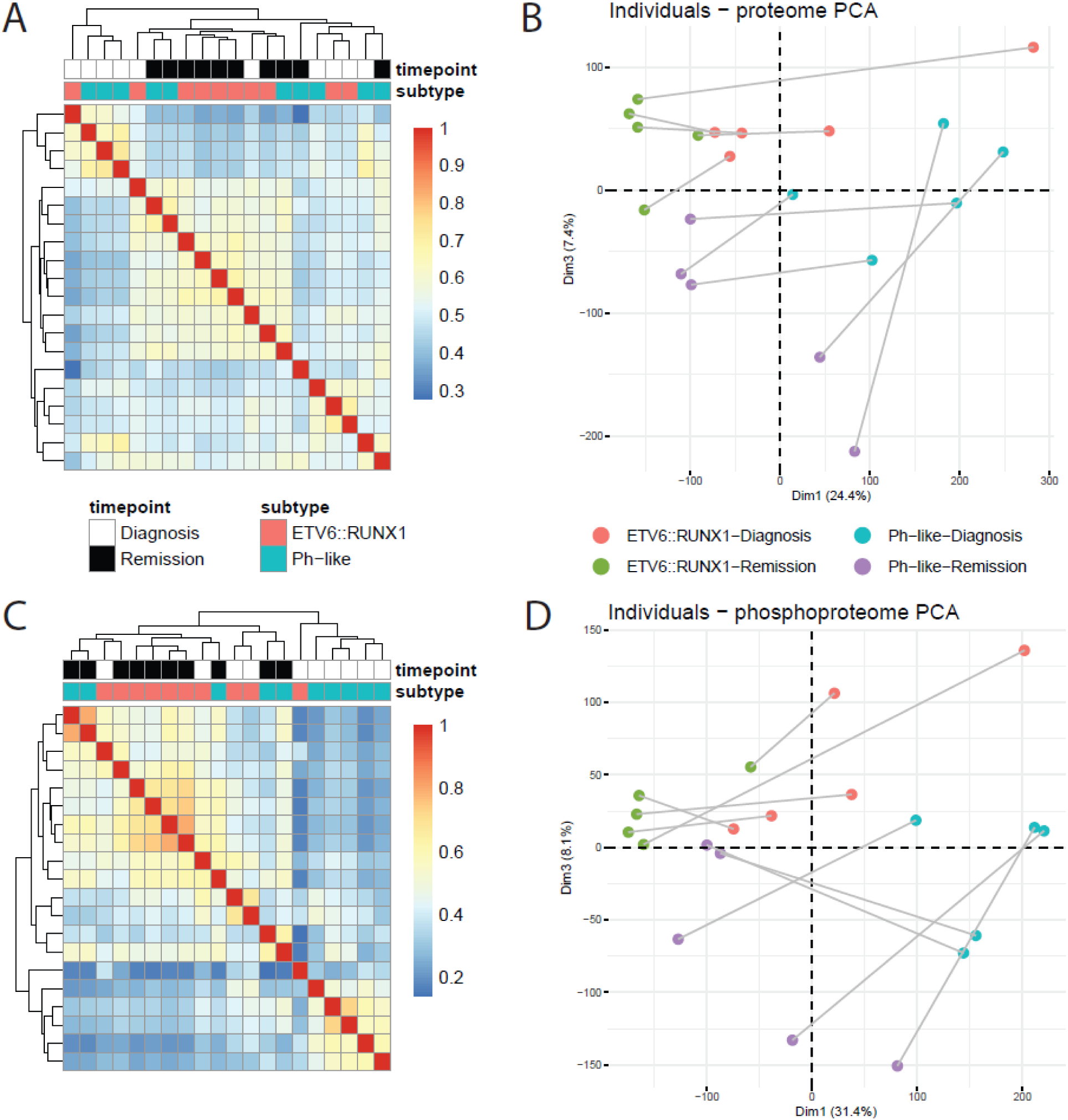
Overall sample similarity across proteomic (A-B) and phosphoproteomic (C-D) assays. A) Pearson correlation matrix of proteomic profiles. B) PCA plot of sample proteomes, with individual patient diagnosis/remission pairs connected. C) Pearson correlation matrix of phosphoproteomic profiles. D) PCA plot of sample phosphoproteomes, with individual patient diagnosis/remission pairs connected.

In contrast to our proteomic profiling where we saw subtype-specific similarity at diagnosis, phosphoproteomic profiling only showed this similarity for Ph-like samples at diagnosis (Figure 2C). As in our proteomic findings, PCA revealed individual patient sample separation between diagnosis and remission (dimension 1) as a large source of variance in the dataset (Figure 2D). Similarly, we did not observe separation based on known characteristics of interest across dimension 2, though again we saw that dimension 3, accounting for similar levels of variance (Supplementary Figure 1C), did show separation according to subtype. In extracting the top five proteins with phosphopeptides contributing to dimensions 1 and 3, we identified phosphosites in ZRANB2, TOP2B, NUMA1, and SPN as drivers of diagnosis vs. remission differences and phosphosites in ARSF1, SRRM1, SUDS3, OCIAD1, and PDHA1 as drivers of subtype-specific separation (Supplementary Figure 1D).

### Differentially expressed protein signatures reveal subtype-specific cancer mechanisms

To identify subtype-specific molecular mechanisms, we performed differential expression analysis between samples collected at diagnosis vs. remission for each subtype. Using cutoffs of minimum 2-fold change in expression and p < 0.05, we identified 495 proteins more highly expressed at diagnosis, and 301 proteins more highly expressed at remission in Ph-like samples (Supplementary Table 1). For *ETV6::RUNX1* samples, we identified 955 proteins more highly expressed at diagnosis and 158 proteins more highly expressed at remission (Supplementary Table 1).

To better understand subtype-specific mechanisms driving leukemias, we looked for overlaps between proteins differentially expressed at diagnosis and remission for each subtype (Figure 3A). We observed that 317 of the proteins were upregulated at diagnosis across both subtypes, suggesting common leukemic mechanisms. Additionally, 577 and 162 proteins were only upregulated at diagnosis in *ETV6::RUNX1* and Ph-like patients, respectively, reflecting subtype-specific proteomic features. Subsequently, we looked for GO, KEGG, and Reactome term enrichment using g:Profiler (Kolberg et al., 2023) to contextualize the differentially expressed proteins. Again, we observe distinct sets of terms enriched for shared and subtype-specific upregulation at diagnosis (Figure 3B; Supplementary Table 2).

**Figure 3.**
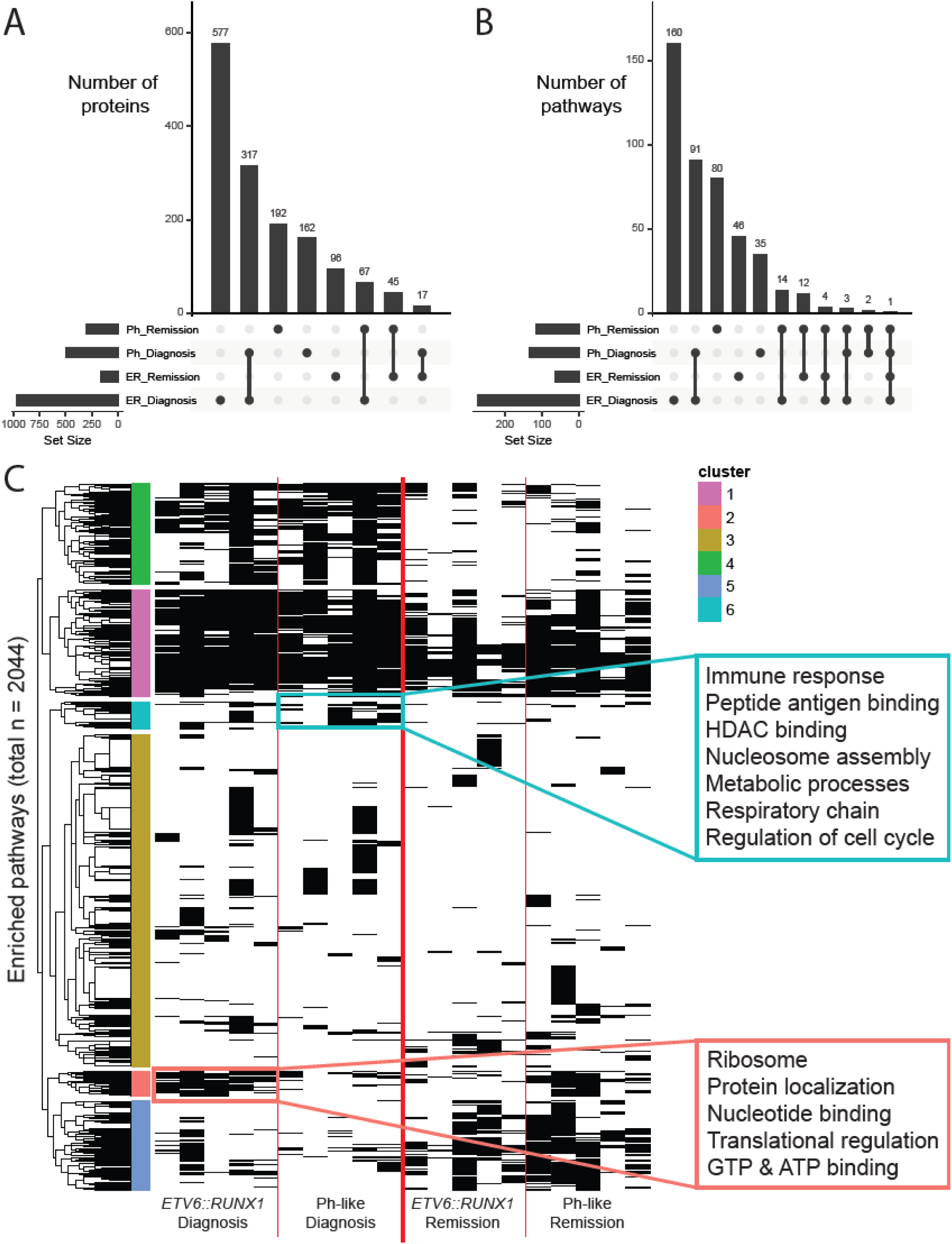
Differential expression of proteins between diagnosis and remission. A) UpSet plot of diagnosis vs. remission differentially expressed proteins for each subtype. B) UpSet plot of pathways enriched in subtype-specific diagnosis vs. remission comparison. C) Pathway enrichment of patient- specific diagnosis vs. remission differentially expressed proteins, focusing on subtype-specific terms enriched at diagnosis.

Enriched at diagnosis for both subtypes, we see terms relating to DNA repair, chromatin remodeling, and transcriptional regulation via RNA Polymerase II. Specific to *ETV6::RUNX1* diagnosis samples, we see terms associated with translational regulation and regulation by p53, an extensively characterized tumor suppressor. Finally, unique to Ph-like diagnosis samples, we see terms associated with cell cycle regulation, specifically metaphase/anaphase transition, as well as SUMOylation.

### Paired diagnosis/remission proteomic comparison reveals common subtype-specific processes

Due to the high degree of heterogeneity observed across the global proteomes of these samples, we leveraged the paired nature of our dataset to explore individual patient comparisons. For each patient, we compared the diagnosis and remission samples, selecting for proteins with at least a 2-fold change in expression between them. For all the proteins more highly expressed in either sample, we looked for enriched pathways to identify mechanistic commonalities.

This analysis identified a total of 2044 pathways enriched in at least one patient sample (Figure 3C). Unsurprisingly, some pathways are enriched across nearly all samples (pink cluster), some enriched at diagnosis regardless of subtype (green cluster), and some enriched at remission regardless of subtype (teal cluster). The majority of pathways are enriched only in a small subset of patient samples, regardless of subtype or timepoint (blue cluster), consistent with the highly individual nature of pediatric leukemias and heterogeneity of global proteomes described in Figure 2.

Of greatest interest to us were the pathways enriched at diagnosis for patients with one subtype, but not the other (yellow and salmon clusters). These pathways correspond to subtype-specific mechanisms driving leukemias. For patients with Ph-like B-ALL, we observe a strong enrichment of terms associated with immune response, chromatin remodeling, metabolic processes, and cell cycle regulation (Supplementary table 3). In contrast, for patients with *ETV6-RUNX1* B-ALL, we primarily observed enrichment of terms associated with ribosomes and translational regulation, as well as nucleic acid binding. These findings suggest that paired diagnosis vs. remission proteomic analysis can identify subtype-specific mechanisms driving pediatric leukemias.

### Individual ‘omics comparisons at diagnosis reveal distinct subtype-specific features

To comprehensively characterize subtype-specific features of the cellular environment, we focused on comparisons at diagnosis between *ETV6::RUNX1* and Ph-like patient samples across three distinct assays: RNAseq, proteomics, and phosphoproteomics. Using existing RNAseq data from patient samples in the CRIB, we identified 11 *ETV6::RUNX1* samples and 6 Ph-like samples (including all patients in Table 1 with the exception of Ph4) with available data. Consistent with prior findings, we observed higher expression of *CRLF2* in Ph-like patient samples (Figure 4A).

**Figure 4.**
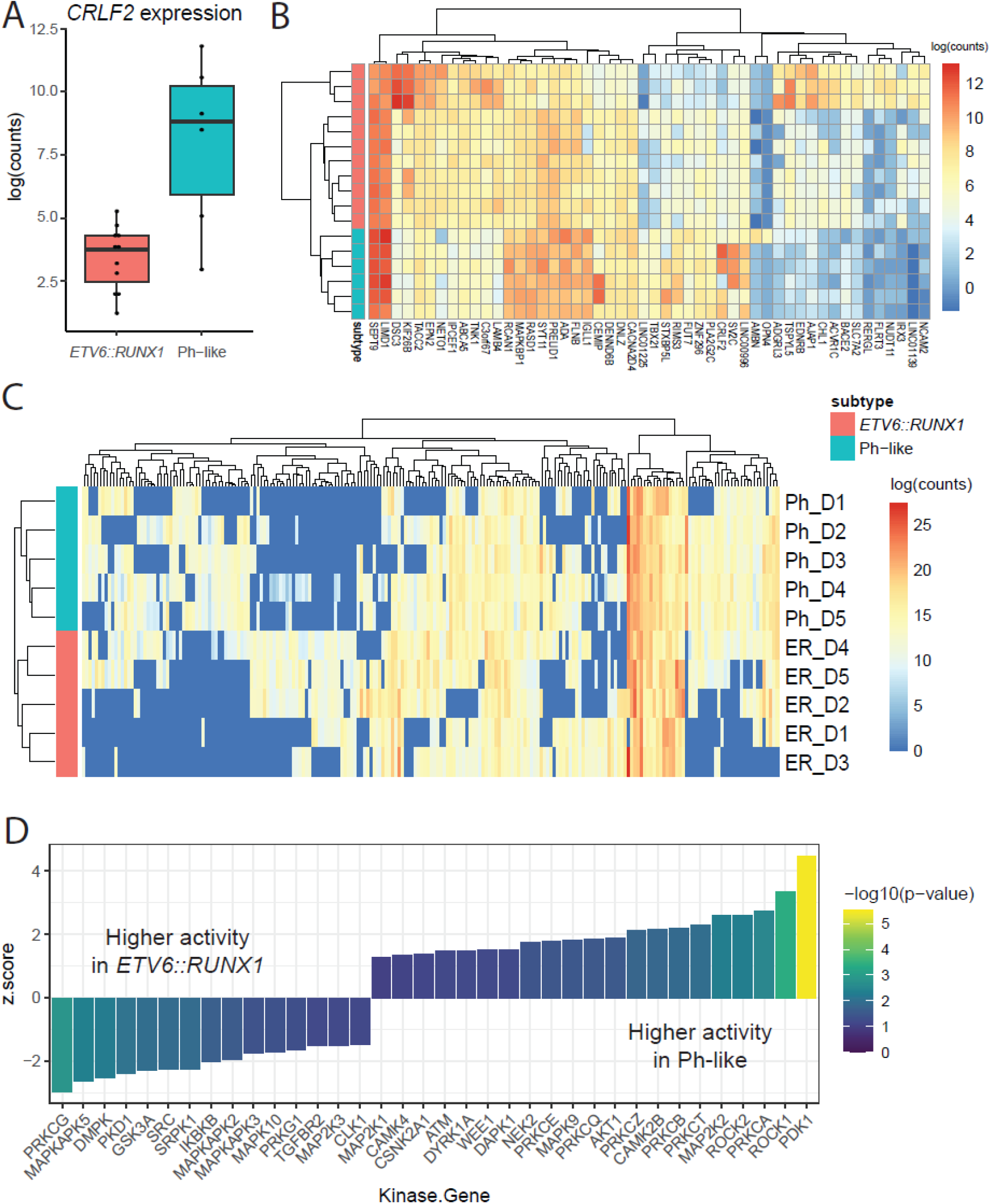
Subtype-specific characteristics at diagnosis. A) Expression of *CRLF2* gene between subtypes. B) Expression heatmap of top 50 genes differentially expressed between subtypes at diagnosis. C) Expression heatmap of 215 proteins differentially expressed between subtypes (detected in at least 4 samples). D) Kinases with statistically significant (p < 0.1) difference in activity, as predicted by KSEA, between Ph-like and *ETV6::RUNX1* samples at diagnosis. Kinases on the right are more active in Ph-like samples, and those on the left are more active in *ETV6::RUNX1* samples.

Differential expression analysis revealed 394 genes more highly expressed in *ETV6::RUNX1* patients and 416 genes more highly expressed in Ph-like patients (p-value < 0.05, Supplementary Table 4). Visualizing the top 50 differentially expressed genes (by p-value), we observed samples clustering according to subtype and distinct gene expression signatures (Figure 4B). While these data recapitulate previous work and identify individual genes of interest, pathway enrichment analysis of these genes did not reveal specific mechanisms of interest, beyond general processes such as development, hematopoietic cell lineage, and differentiation (Supplementary Table 5).

Comparing proteomic profiles across the subtypes at diagnosis using cutoffs of minimum 2-fold change in expression and p < 0.05, we identified 192 proteins more highly expressed in Ph-like samples and 355 proteins more highly expressed in *ETV6::RUNX1* samples (Supplementary Table 1). Visualizing expression of differentially expressed proteins at diagnosis, we saw samples clustering according to subtype, as well as distinct sets of proteins whose expression was primarily detected only in one subtype (Figure 4C). Pathway enrichment analysis using g:Profiler (Kolberg et al., 2023) showed a strong bias toward rRNA, ribosomal, and translational processes in *ETV6::RUNX1* samples. In contrast, Ph-like samples showed enrichment for processes involved in ATP-dependent regulation and calcium-dependent signaling (Supplementary Table 2).

To identify processes of interest from our phosphoproteomic profiling, we used the KSEA App (Wiredja et al., 2017) to predict which kinases showed differences in activity levels between Ph-like and *ETV6::RUNX1* samples at diagnosis, based on differences in phosphopeptide abundance. This analysis identified a total of 36 kinases whose activity was significantly different (p < 0.1 to expand the search space) between the two subtypes (Figure 4D; Supplementary Table 6). In Ph-like samples, we saw elevated activity of kinases including PDK1, ROCK1, AKT1, CAMK2B, and numerous members of the Protein Kinase C (PKC) family including PRKCA, PRKCT, and PRKCB, which are known to be activated by calcium signaling. In contrast, in *ETV6::RUNX1* patient samples, we saw higher activity of PRKCG, DMPK, and multiple members of MAPK signaling pathways, including MAPKAPK5, MAPKAPK2, MAPKAPK3, and MAPK10.

It is notable that we observed consistency between our proteomic finding of increased calcium-dependent signaling in Ph-like samples and our phosphoproteomic finding of PKC family activity in these samples, though there was no evidence of these subtype-specific processes gleaned from RNAseq analysis. These congruent findings support the utility and value of proteomic and phosphoproteomic methods to better understand subtype-specific molecular mechanisms in pediatric leukemias.

### Multi-omic analysis reveals subtype-specific regulatory relationships

To explore how a combination of all three datasets informs sample similarity, we performed multiple co-inertia analysis (MCIA) using the omicade4 package (Meng et al., 2014). Upon performing MCIA, we see clear separation of samples according to subtype across dimension 2 (Figure 5A). Interestingly, dimension 1, which accounts for the largest variance in the dataset, appears primarily to separate sample ER4 from other samples. This is especially striking as this is the only *ETV6::RUNX1* sample (ER4) where the “second hit” driving leukemogenesis is a partial loss of *PAX5* rather than loss of *ETV6*, supportive of a distinct molecular mechanism that has been observed in this recently described subtype of *ETV6::RUNX1* B-ALL (Z. Li et al., 2025).

**Figure 5.**
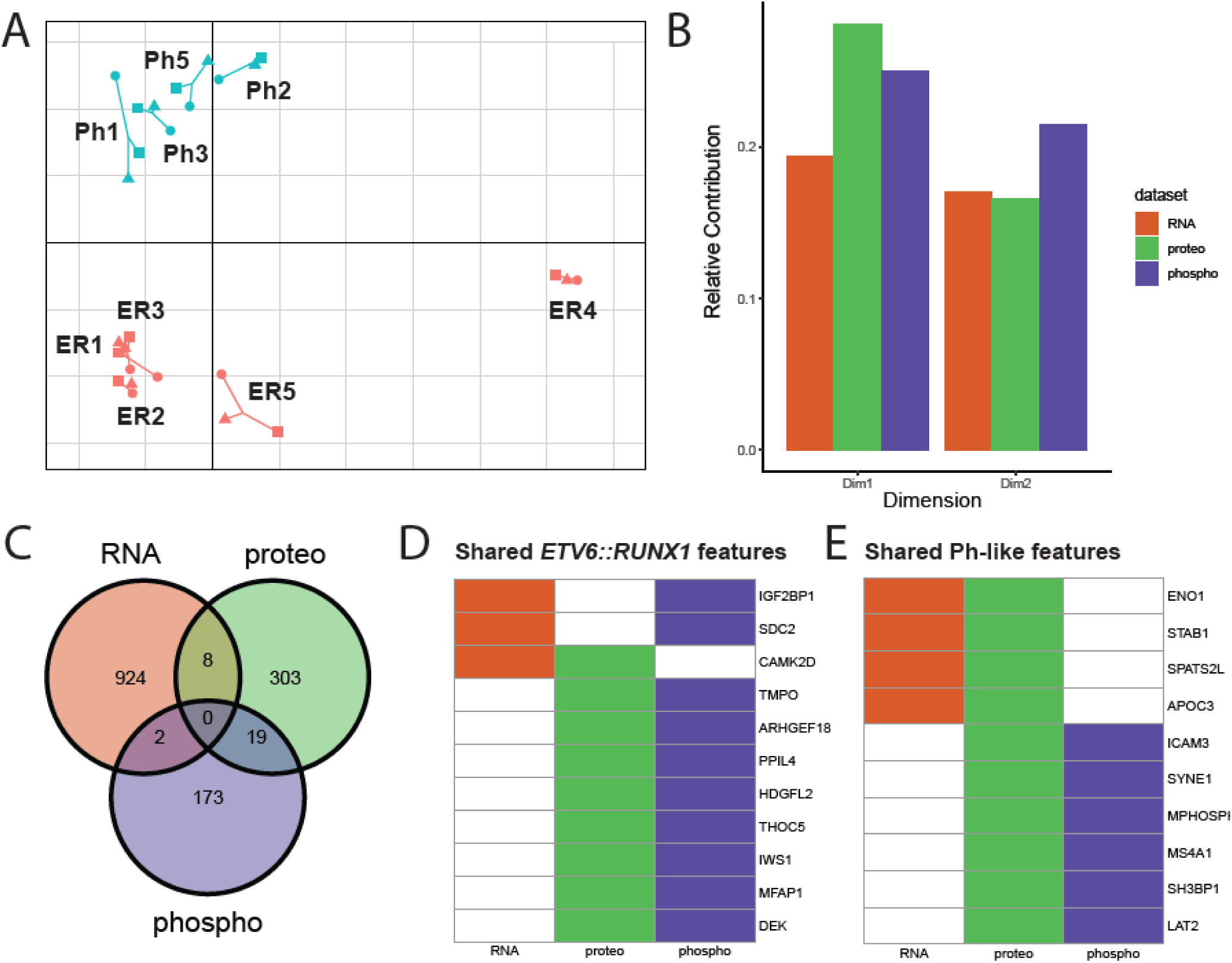
Multiple Co-Inertia Analysis (MCIA) of Ph-like vs. *ETV6::RUNX1* samples at diagnosis. A) MCIA plot dimensions 1 (x-axis) and 2 (y-axis) showing sample similarity using all three modalities. B) Relative contribution (based on pseudo-eigenvalues) of each dataset to dimensions 1 and 2 of MCIA plot. C) Overlap of dimension 2 subtype-biased features (|Dim2| > 1), which separate subtypes across modalities. D-E) All drivers that came up in multiple datasets toward each subtype, black = putative subtype-biased feature in this dataset, white = not considered a subtype-biased feature in this dataset.

We also queried the pseudo-eigenvalue space of the MCIA result to determine the relative contributions of each assay to variance across dimensions 1-2 (Figure 5B). Proteomics followed by phosphoproteomics contributed more to the separation of *ETV6::RUNX1* sample ER4 from others across dimension 1, while phosphoproteomics contributed the most to separation of the subtypes across dimension 2. It is especially interesting to note that although the RNAseq assay captured nearly an order of magnitude more features (genes, proteins, or phosphopeptides) than the other two assays, it did not contribute as much to the overall variance of the sample space.

Since we saw clear separation of subtypes across dimension 2, we leveraged the feature space to identify subtype-specific patterns across datasets. We selected features across all three assays that had highly positive or negative weights for this dimension, identifying 942 genes, 329 proteins, and 216 phosphopeptides of interest, or subtype-biased features. Mapping each feature back to the corresponding gene name, we were surprised to find that very few features appeared to be drivers across multiple datasets (Figure 5C).

Putative *ETV6::RUNX1*-biased features shared across datasets include IGF2BP1, SDC2, TMPO, and TMEM40 (Figure 5D). Putative shared Ph-like-biased features include ENO1, ICAM3, SH3BP1, and LAT2 (Figure 5E). Although they may not have been directly linked to pediatric B-ALL, many of the shared features identified through this analysis have been implicated in other cancers, predicted to act as tumor suppressors, involved in cell signaling, or associated with regulation of hematopoiesis.

### Features identified through MCIA suggest novel subtype-specific regulatory processes

Further investigation into top features of interest, both those shared across datasets and not, revealed a wide array of regulatory patterns between RNAseq, proteomics, and phosphoproteomics (Figure 6A-C, respectively). Some showed consistent patterns across the three assays – for example *IGF2BP1* gene expression, IGF2BP1 protein expression, and phosphorylation of IGF2BP1-S181 were all higher in *ETV6::RUNX1* samples. Other drivers showed discordant trends – for example *CHD3* gene expression was higher in Ph-like samples, CHD3 protein expression was higher in *ETV6::RUNX1* samples, and phosphorylation of CHD3 at S1660 and S1664 was higher in Ph-like samples. And still other proteins, including BCLAF1, showed increased phosphorylation at some sites and decreased phosphorylation at others, suggesting regulation by multiple kinases with potentially disparate effects on downstream targets.

**Figure 6.**
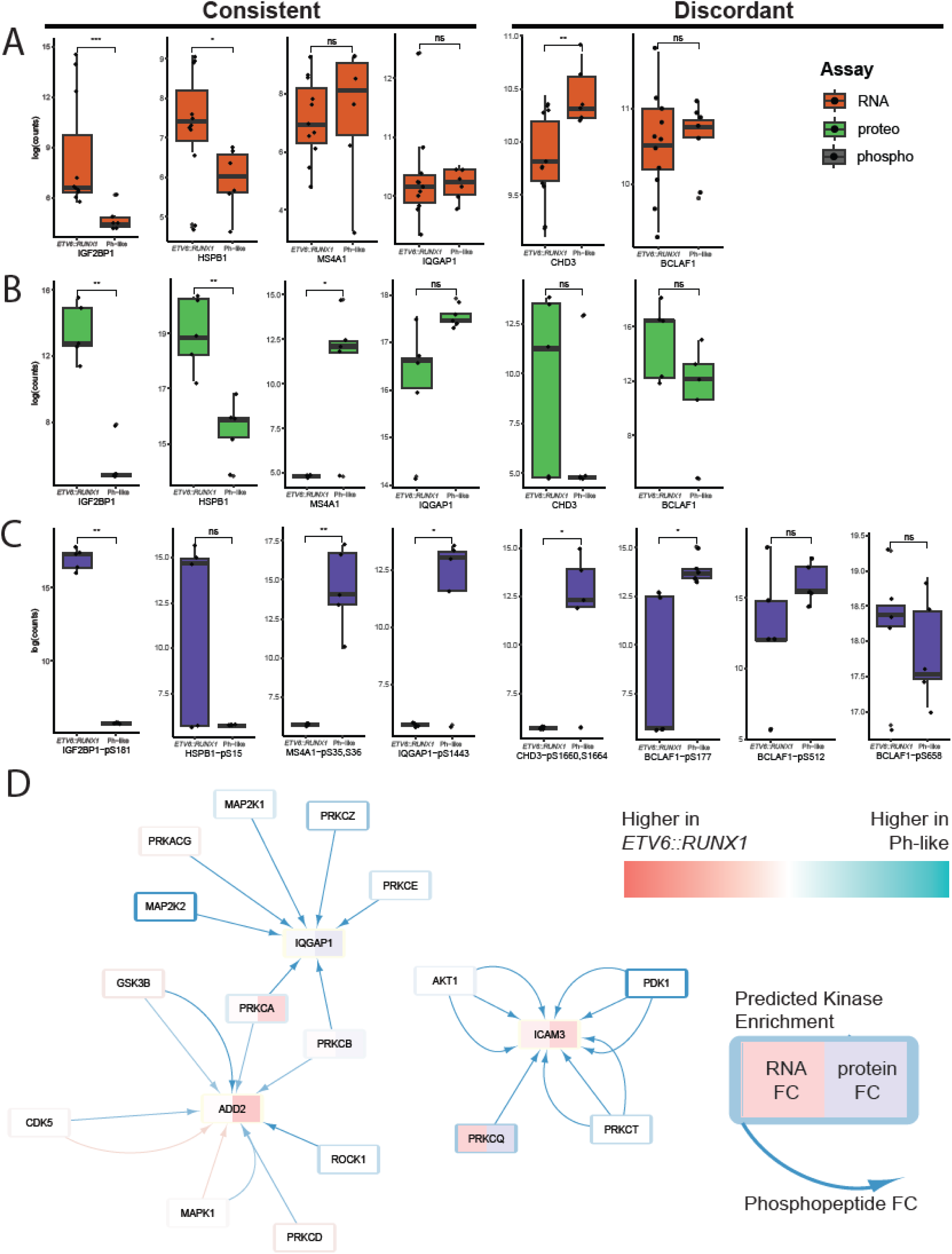
Subtype-specific mechanisms identified through multi-omic analysis. A-C) Expression of selected drivers in RNA (A), proteomic (B), and phosphoproteomic (C) datasets. ns = not significant, * = p < 0.05, ** = p < 0.01, *** = p < 0.001. D) Network visualization of PKC-related activity incorporating fold change data from RNA, proteomic, and phosphoproteomic assays. Each arrow represents phosphorylation of an individual site on the target protein. White reflects log2FC = 0 or no data captured for the assay.

To visualize the connections between datasets, we leveraged the kinase target data from our phosphoproteomic analysis using the KSEA App to build a network in Cytoscape (Supplementary Table 7). Treating each phosphosite as a unique phosphorylation event, we were able to visualize connections between proteins and identify sub-networks of interest. Focusing on calcium-dependent signaling processes, which we saw enriched in Ph-like samples across proteomic and phosphoproteomic data, we were able to identify interactions with proteins implicated in actin cytoskeleton remodeling (Figure 6D). Significantly, these interactions appeared to be largely driven by kinase activity to a far greater extent than differences in protein or RNA abundance between the two subtypes.

This work demonstrates the value of an integrative analysis such as MCIA to combine transcriptomic, proteomic, and phosphoproteomic data for the study of pediatric leukemias. Through this analysis, we identified subtype-specific characteristics both shared across datasets, and showing distinct expression patterns. Exploring such patterns in larger-scale studies will lay the groundwork for developing mechanistic hypotheses about subtype-specific processes in leukemogenesis and treatment resistance.

## Discussion

Characterization of leukemias across multiple cellular facets – genomic, transcriptomic, proteomic, and phosphoproteomic – facilitates identification of subtype-specific features and molecular mechanisms (Vllahu et al., 2025). Data integration across these modalities enables the identification of key regulatory interactions driving phenotypes and targetable signaling cascades. Although proteomic profiling is becoming more common in the study of pediatric leukemia (Kourti et al., 2023; López Villar et al., 2014), most studies to date have been limited by either a focus on cell lines or a lack of integration with other ‘omics data. This study is one of the first to integrate genomic, transcriptomic, proteomic, and phosphoproteomic data generated from primary pediatric leukemia patient samples. Our comprehensive profiling of two well-characterized subtypes of pediatric B-ALL, Ph-like (*BCR::ABL1*-like) and *ETV6::RUNX1*, identifies features of interest as well as subtype-specific regulatory interactions.

We performed comparative analysis of proteomic and phosphoproteomic data between matched patient samples at diagnosis and remission. This enabled us to identify salient features that are both common across all patients and restricted to a particular subtype. *ETV6::RUNX1* B-ALL showed a high degree of translational regulation, as well as activation of MTOR and MAPK signaling pathways. In contrast, in Ph-like B-ALL we observed increased calcium-dependent signaling, and increased activity of CAMK and AKT-driven signaling. In addition to standard differential expression analysis, we wanted to leverage the paired nature of our data and performed individual patient diagnosis vs. remission comparison, similar to clinical tumor vs. normal approaches. This analysis also identified translational regulatory processes as enriched in *ETV6::RUNX1* patient samples at diagnosis and cell cycle regulation enriched in Ph-like samples at diagnosis, though interestingly did not capture calcium-dependent signaling in this context (Figure 3C). We were encouraged to see some consistency between the two approaches, though unsurprisingly also observed discrepancies, which were likely amplified due to the lack of statistical power in individual patient diagnosis vs. remission analysis.

As we were particularly interested in the power of multi-omic analysis, we performed single-assay comparisons between the two subtypes. Transcriptional comparison successfully identified genes associated with the Ph-like subtype (Harvey et al., 2013; Reshmi et al., 2017). To date, most proteomic studies of pediatric ALL patient samples have focused on comparison of high-risk vs. low-risk patients, as opposed to subtype-specific comparison (Braoudaki et al., 2013; Xu et al., 2017). Notably, a recent study by Gupta et al., performed global proteomic characterization of Ph-like B-ALL patient samples (Gupta et al., 2023). Surprisingly, we observed limited overlap with their findings – of their top 24 proteins overexpressed in Ph-like samples (compared to non-Ph-like samples), none were statistically significant in our comparison at diagnosis.

Interestingly, we did observe that some of these proteins of interest, including JCHAIN and DDX27, were statistically significantly more highly expressed in Ph-like samples at diagnosis than at remission, while others, including NDUFA2, IGHA1, and CHMP4B, were more highly expressed at remission in Ph-like samples than *ETV6::RUNX1* samples, though not at diagnosis (Supplementary Table 1). These differences may be due to our specific selection of two subtypes for comparison versus their comparison of Ph-like to “non-Ph-like” cases which may have a variety of drivers and underlying mechanisms. Additionally, it is important to keep in mind that pediatric cancers often behave differently from their adult counterparts, which may explain discrepancies between our pediatric study and their adult focus (Ahmed et al., 2018; Milan et al., 2019).

Pathway analysis of our proteomic results suggested a role for increased calcium-dependent signaling in Ph-like samples (Supplementary Table 2). This is supported in our phosphoproteomic results with predicted increased activity of CAMK2B and members of the PKC signaling family in Ph- like samples (Figure 4D). These observations support previously documented findings that calcium signaling, and PKC activity specifically, may play a vital role in the development and progression of leukemias (Immanuel et al., 2022; Redig & Platanias, 2008). Interestingly, atypical protein kinase C λ/ι has been specifically implicated in leukemic transformation of *BCR::ABL1*+ cells (Nayak et al., 2019).

Some of the best characterized hallmarks of Ph-like leukemia include upregulation of CRLF2 activity and JAK/STAT signaling (Harvey et al., 2010; Roberts et al., 2012; Schmäh et al., 2017). While we did observe higher *CRLF2* gene expression (Figure 4A), our proteomic analysis did not capture CRLF2 or JAK/STAT family proteins. Similarly, the SMOAC method enrichment approach (Choi et al., 2017) for phosphoproteomics selects for phospho-serines and phospho-threonines, thus limiting our ability to observe direct effects of tyrosine kinase activation that have previously been described in Ph- like B-ALL. Future efforts should leverage more sensitive proteomic methods and phospho-tyrosine selecting phosphoproteomic approaches to confirm and extend these findings. Nonetheless, our approach has identified functional subtype-specific signaling cascades which may open avenues for non-TKI therapeutic targeting.

Most significantly, our integrated analysis of transcriptomic, proteomic, and phosphoproteomic data facilitated a rapid, functionally focused analysis across data modalities. Based on clinical characterization of these patient samples (Table 1), we see that not only do the Ph-like patients in this study have diverse drivers, as is common for the subtype (Roberts et al., 2014b; van Outersterp et al., 2024), but the *ETV6::RUNX1* patients also show diverse “second hit” drivers beyond the *ETV6::RUNX1* rearrangement itself (Kaczmarska et al., 2023; Z. Li et al., 2025; Rodríguez-Hernández et al., 2021). Strikingly, our multi-omic analysis clearly picked up the difference, as evidenced by the clear separation of patient sample ER4 from other *ETV6::RUNX1* samples across Dimension 1 of the MCIA (Figure 5A). It is also interesting to see that this separation was more strongly driven by proteomic and phosphoproteomic than RNA data (Figure 5B), supporting the idea that proteomic and phosphoproteomic profiles show a more comprehensive functional readout of the cell state than the transcriptome.

This integrated approach also creates a foundation for developing mechanistic hypotheses about subtype-specific regulatory interactions. We were able to quickly identify features that were consistently more highly expressed in one subtype across all assays (Figure 5 C-E), including a bias for *MS4A1* gene expression, MS4A1 protein expression, and MS4A1-S35,S36 phosphorylation in Ph-like cases. The MS4A1 protein, also known as CD20, has been shown to be upregulated, and targetable, in chronic lymphocytic leukemia (Pavlasova et al., 2016) and expressed in leukemic stem cells in ALL (Blatt et al., 2018). Similarly, we were able to directly query the dataset for features showing opposing trends across assays (i.e. higher protein expression in one subtype but higher phosphorylation in the other) to identify features of interest such as CHD3 (Figure 6A-C). Although CHD3 has not previously been associated with leukemias, it is a component of the NuRD complex which acts as a chromatin remodeler and is involved in the regulation of numerous downstream targets (Denslow & Wade, 2007; Hoffmeister et al., 2017). The discordant expression pattern across assays suggests that there may be tight post-translational regulation of this protein, and its involvement in specific subtypes of pediatric B- ALL merits further investigation.

Finally, leveraging the kinase target data we obtained by running the KSEA App for kinase activity prediction on our phosphoproteomic comparison of the two subtypes at diagnosis, we were able to visualize interactions between proteins and patterns across the data modalities. We were able to identify several sub-networks of calcium-dependent kinases phosphorylating targets involved in actin cytoskeleton remodeling (Figure 6D). The connection to the cytoskeleton is particularly interesting as BCR-ABL kinase activity has been shown to regulate the actin cytoskeleton in Ph+ leukemic cells, a process which has been suggested to play a role in the maintenance of leukemic stem cells as well as therapeutic resistance in cells expressing the fusion protein (McWhirter & Wang, 1993; Preisinger & Kolch, 2010; Wang et al., 2007). Although this has not been described in Ph-like leukemias to date, possibly due in part to the abundance of distinct drivers for this subtype, it presents an exciting avenue for further investigation.

While further work with larger sample sizes will be required to validate and extend these findings, this study represents one of the first efforts toward comprehensive multi-omic characterization of specific pediatric leukemia subtypes. Taken together, these data demonstrate the utility of integrated multi-omic analysis in identifying subtype-specific regulatory mechanisms. Application of this approach will facilitate the development of novel mechanistic hypotheses which will in turn lead to improved stratification and treatment selection for children diagnosed with B-ALL.

## Acknowledgements

We would like to acknowledge the Children’s Mercy Research Institute Biorepository (CRIB) for assistance with sample collection and processing. This work was supported by an MCA Partners Advisory Board grant from Children’s Mercy Hospital (CMH) and The University of Kansas Cancer Center (KUCC). The proteomics data were collected in the Mass Spectrometry and Proteomics Core facility utilizing the Orbitrap Ascend Tribrid System that was purchased with funds provided by the University of Kansas Cancer Center, which is supported by the National Cancer Institute Cancer Center Support Grant P30 CA168524.

---

**Supplementary Figure 1.**
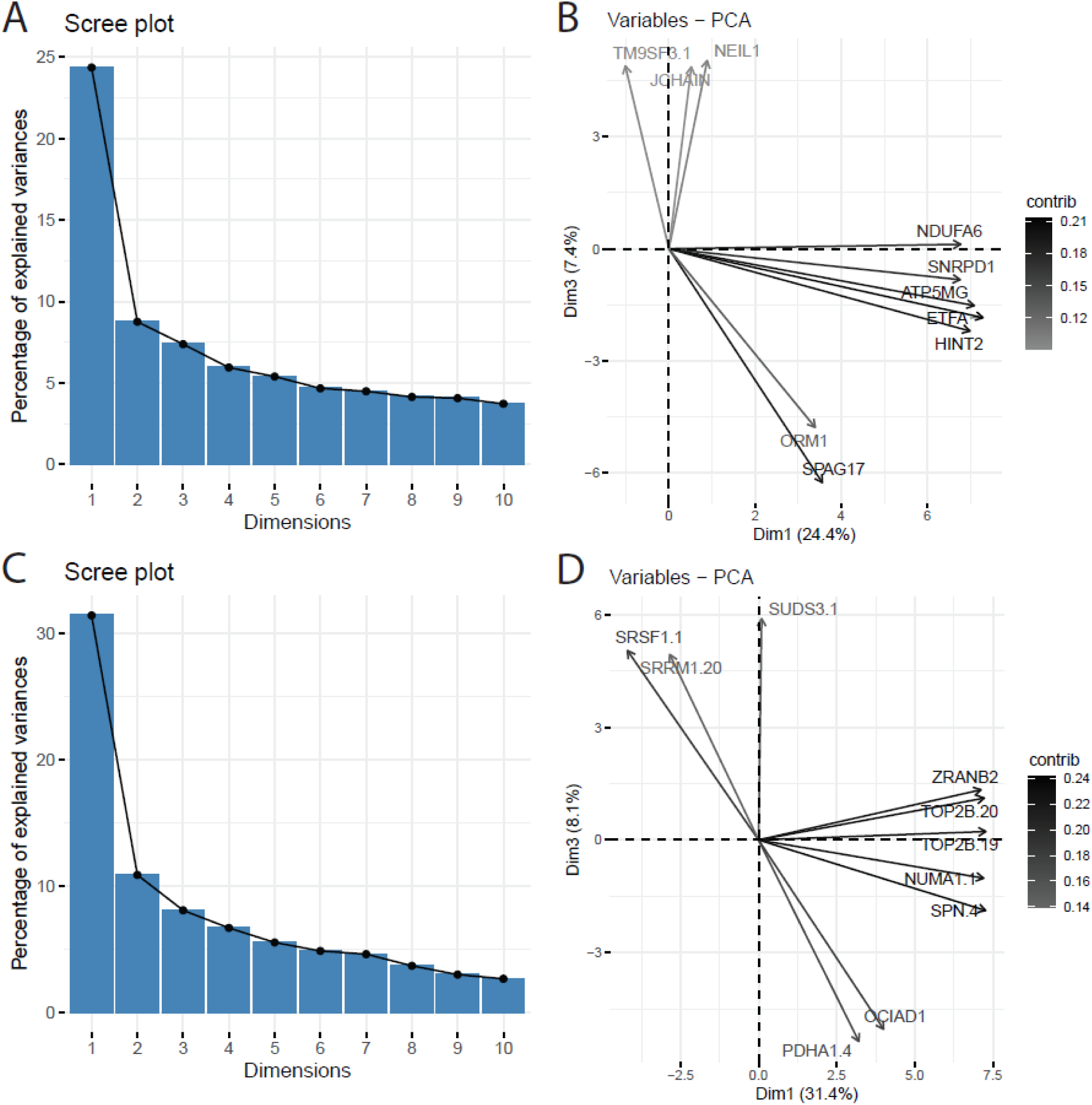
Scree plots and top 5 variables driving dimensions 1 and 3 for proteomic (A-B) and phosphoproteomic (C-D) PCAs.

## Supplementary Table legends

**Supplementary Table 1**. Proteins statistically significantly differentially expressed (p < 0.05, |fold change| > 2) across comparisons (including directionality). 1 = statistically significant, 0 = not statistically significant. ER = *ETV6::RUNX1*, Ph = Ph-like, D = diagnosis, N = remission. For comparisons between subtypes, column names are [timepoint]_PhvER_[Ph/ER – subtype where higher]. For comparisons between timepoints, column names are [subtype]_DvN_[D/N – timepoint where higher].

**Supplementary Table 2.** Pathway enrichment results on differentially expressed proteins using gProfiler. The 8 sets of results correspond to the 8 directional lists of differentially expressed proteins in Supplementary Table 1 and are in the same order.

**Supplementary Table 3.** Pathway enrichment results on differentially expressed proteins based on individual patient diagnosis vs. remission comparisons. Columns labeled according to patient identifier, with D = enriched at diagnosis and N = enriched at remission (p < 0.05).

**Supplementary Table 4.** Statistically significant (p < 0.05) genes differentially expressed between Ph- like and *ETV6::RUNX1* patient samples at diagnosis. Positive log2FC = higher in Ph-like, negative log2FC = higher in *ETV6::RUNX1*.

**Supplementary Table 5.** Pathway enrichment results on differentially expressed genes between Ph- like and *ETV6::RUNX1* patient samples at diagnosis. Comparison 1 corresponds to genes more highly expressed in Ph-like samples, comparison 2 corresponds to genes more highly expressed in *ETV6::RUNX1* samples.

**Supplementary Table 6.** Differential kinase activity between Ph-like and *ETV6::RUNX1* patient samples at diagnosis, as predicted by KSEA App. Positive enrichment/z-score = higher in Ph-like, negative enrichment/z-score = higher in *ETV6::RUNX1*.

**Supplementary Table 7.** Differentially expressed phosphopeptide targets (substrates) between Ph-like and *ETV6::RUNX1* patient samples at diagnosis, generated by KSEA App. Positive log2FC = higher in Ph-like, negative log2FC = higher in *ETV6::RUNX1*.

